# White-tailed deer milk exhibits SARS-CoV-2 neutralizing antibodies and synergistic mechanisms that contribute to rapid viral RNA degradation

**DOI:** 10.64898/2026.02.11.705256

**Authors:** Lorena Tobar, Wraylyn Smith, Hannah A. Raso, Katherine L. Brown, Alessandro Ceci, Matthew G. Urbano, Clinton Roby, Dzenis Mahmutovic, Andrew P. Biesemier, Daniel G.S. Capelluto, Ignacio Aiello, Dominic S. Raso, Carla V. Finkielstein

**Affiliations:** Fralin Biomedical Research Institute at VTC, Roanoke, VA 24016; Department of Dairy Science, Virginia Tech, Blacksburg, VA 24060; Pathology Consultants of Central Virginia, Lynchburg, VA 24501; Department of Pathology, University of Virginia, Charlottesville, VA 22908; Department of Biological Sciences, Virginia Tech, Blacksburg, VA 24060; Fralin Life Sciences Institute, Blacksburg, VA 24060

**Keywords:** SARS-CoV-2, white-tailed deer, COVID-19, milk, immunity, virus stability

## Abstract

White-tailed deer (WTD) represent the most significant SARS-CoV-2 wildlife reservoir in North America, yet the role of antiviral mechanisms in vertical transmission remains unexplored. We investigated SARS-CoV-2 antibody responses and viral stability in milk from lactating WTD and humans to characterize species-specific antiviral mechanisms. SARS-CoV-2 neutralizing antibodies were detected in milk and serum in WTD specimens using complementary immunoassays, providing the first evidence of humoral immune responses in wildlife milk. Despite antibody presence indicating prior SARS-CoV-2 exposure, viral RNA was undetectable in all WTD milk samples. This pattern aligns with observations in human milk, where viral RNA was also undetectable both during active infection (when nasal swabs were positive) and during antibody-positive periods following recovery. *In vitro* stability studies revealed striking species differences: all SARS-CoV-2 variants (A, B.1.1.7, BA.1.1.529) rapidly degraded in WTD milk within 30 min at physiological temperatures, while remaining mostly stable in human milk for up to 60 min. Biochemical characterization identified multifactorial degradation mechanisms in WTD milk, including 5-20 fold elevated mineral concentrations (sodium, magnesium, phosphorus, and potassium), enhanced protease activity, and increased lactoperoxidase levels. Individual mineral supplementation revealed variant-specific susceptibilities, with B.1.1.7 showing pronounced sensitivity to ionic stress. Mechanistic studies demonstrated synergistic effects between elevated ionic concentrations and proteolytic activity, with heat-labile and heat-stable degradation pathways contributing to viral inactivation. These findings reveal that WTD milk possesses intrinsic antiviral properties fundamentally different from human milk, representing an evolutionary adaptation that may impact viral persistence and transmission dynamics in wildlife populations. These findings reveal antiviral mechanisms in WTD milk that represent a previously unrecognized component of pathogen control in wildlife reservoirs, with important implications for understanding wildlife-pathogen interactions and zoonotic risk assessment.

**Author Summary:** White-tailed deer (WTD) have become the primary wildlife reservoir for SARS-CoV-2, with millions of infected animals across North America. Despite this significance, the presence of protective antibodies and viral behavior in WTD milk remained unexplored. We collected milk samples from lactating WTD during hunting seasons to investigate whether WTD produce neutralizing antibodies similar to those found in human milk and to examine how the virus behaves in this biological fluid.

Our analysis revealed that WTD milk contains antibodies capable of neutralizing SARS-CoV-2. When we compared viral stability between WTD and human milk, we observed that WTD milk rapidly degrades viral genetic material within 30-40 min, while the same virus remains stable in human milk for over an hour. We identified that WTD milk contains mineral concentrations 5-20 times higher than human milk, including elevated levels of sodium, magnesium, and potassium, along with enhanced enzyme activity that breaks down viral components.

These findings indicate that WTD milk functions as a protective barrier rather than a transmission route. This has implications for understanding viral persistence in wildlife populations and assessing potential risks to human health. Our work demonstrates that deer milk possesses multiple biological defense mechanisms that may protect offspring from viral infections, contributing to our understanding of wildlife immunity and pandemic preparedness.

## Introduction

The severe acute respiratory syndrome coronavirus 2 (SARS-CoV-2) pandemic has demonstrated extensive cross-species transmission potential, with natural infections documented in over 30 mammalian species through both human-to-animal and animal-to-animal transmission pathways [1–4]. Among wildlife species, white-tailed deer (*Odocoileus virginianus*, hereafter WTD) have emerged as the most significant reservoir since its initial detection in late 2021 followed by a widespread infection across North America [5–7]. SARS-CoV-2 RNA and seroprevalence studies report infection rates of 15-40% across surveyed populations, with evidence of sustained viral circulation and adaptation within WTD populations [6–9]. In some localities, prevalence rates were even higher, underscoring a strong heterogeneous spatial distribution of infection [5, 8, 10]. The epidemiological significance of this reservoir is amplified by the estimated 30-34 million WTD population across North America and the frequent human-wildlife interfaces that occur through hunting, farming, and suburban contact [11].

The establishment of wildlife reservoirs complicates pandemic control efforts and creates ongoing risks for viral persistence, evolution, and potential reverse zoonotic transmission [12]. Understanding viral transmission dynamics in these reservoir populations involves investigating pathogen behavior in biological matrices that facilitate maternal-offspring transmission, particularly milk. Research on SARS-CoV-2 in human milk shows that viral RNA is rarely detected, the infectious virus is absent, and no case of infant infection has been linked to breast milk transmission [13, 14]. However, recent studies have demonstrated that SARS-CoV-2 components can persist in breast tissue in select long COVID cases, with viral nucleocapsid and spike proteins plus viral RNA detected months after acute infection, colocalizing with macrophages in mammary tissue [15, 16]. This tissue-level viral persistence contrasts with the consistent absence of infectious virus in breast milk, indicating that mammary glands may serve as viral reservoirs without facilitating milk-borne transmission. Conversely, neutralizing antibodies, mainly IgA, are consistently detected in human milk from infected and vaccinated mothers, suggesting passive immune protection for nursing infants [17].

Despite these findings in human milk, no studies have investigated SARS-CoV-2 dynamics in milk from infected wildlife populations. This knowledge gap is particularly significant as human and other mammalian milks share a common functional foundation but exhibit vast, phylogenetically and ecologically linked differences in macronutrients, micronutrients, bioactive compounds, pH, ionic strength, and antimicrobial factors across species [18, 19], which could markedly influence viral stability and immune factor activity compared to human milk. For example, human milk is characterized by relatively low protein and fat content but high lactose concentrations, whereas ruminant species such as sheep and buffalo produce milk substantially richer in solids and fat [20]. Mineral profiles vary dramatically between species, with ruminant milks generally containing elevated calcium and magnesium levels compared to human milk, while some species like camels and goats show distinctive potassium or iron concentrations [19, 21]. Among cervids, including white-tailed deer (WTD), milk demonstrates unusually high protein, fat, calcium, and distinctive fatty acid profiles including elevated branched-chain fatty acids compared to domestic dairy ruminants [21–23]. Additionally, species-specific differences in antimicrobial factors, lipid composition, and bioactive compounds create distinct biochemical environments that could substantially affect viral persistence and immune factor activity in ways that cannot be predicted from human milk studies alone [24–26].

The protective environment observed in human milk, characterized by minimal viral transmission and abundant neutralizing antibodies, cannot be assumed to exist in wildlife species. This assumption is particularly important given that other pathogens demonstrate variable stability in ruminant milk, with viruses like Avian influenza A (H5N1) [27] and Rift Valley fever showing transmission potential through unpasteurized raw milk, underscoring the pathogen-specific nature of milk-based transmission risk [28–30]. Understanding species-specific viral dynamics is critical for both ecological and public health assessment. Ecologically, milk-transmitted maternal immunity affects population-level susceptibility patterns in wildlife reservoirs [31, 32], while from a public health perspective, even brief viral persistence in deer milk could influence transmission dynamics during production, processing, or consumption of WTD-derived products [33].

Given the established differences in milk composition between species and our histological findings that WTD mammary tissue contains rare SARS-CoV-2-positive histiocytes that could serve as a potential source of cell-associated viral particles in milk at very low levels, we determined the stability of SARS-CoV-2 and characterized the biochemical environment of WTD versus human milk to identify factors influencing viral persistence. Our results demonstrate the presence of neutralizing antibodies and identify distinct physicochemical properties in WTD milk that promote rapid viral RNA degradation, providing the first characterization of SARS-CoV-2 dynamics in wildlife milk and revealing mechanisms that may explain the apparent viral clearance observed in field samples.

## Results

### White-tailed deer and human specimen collection during regional SARS-CoV-2 variant transitions

Lactating white-tailed deer (WTD) samples were collected from central Virginia (Lynchburg area), USA, during the 2022-2024 hunting seasons (Figure S1). In the same geographic area and timeframe, SARS-CoV-2 variant distribution data were obtained from 14,446 human swab samples collected using our previously established protocols to establish regional viral context [34]. Of these, 654 RT-qPCR-positive specimens underwent whole-genome sequencing and variant classification, with Pangolin lineages grouped into four categories for analysis: Omicron, XBB, JN.1, and other variants (Figure 1A). Sample collection timepoints for WTD (orange bars) and human subjects h.s.#1 and h.s.#2 (yellow and purple bars) were mapped against prevalent regional variants to establish temporal context (Figure 1A, WTD, h.s.#1 and h.s.#2 lines). This temporal mapping confirmed alignment between individual infections and regional circulation patterns, with whole genome sequencing and viral characterization identifying EG.5.1.1 (XBB lineage) in h.s.#1 (8/31/2023) and BA.5 in h.s.#2 (08/2022), consistent with the respective periods of regional variant dominance (Figure 1A).

**Figure 1.**
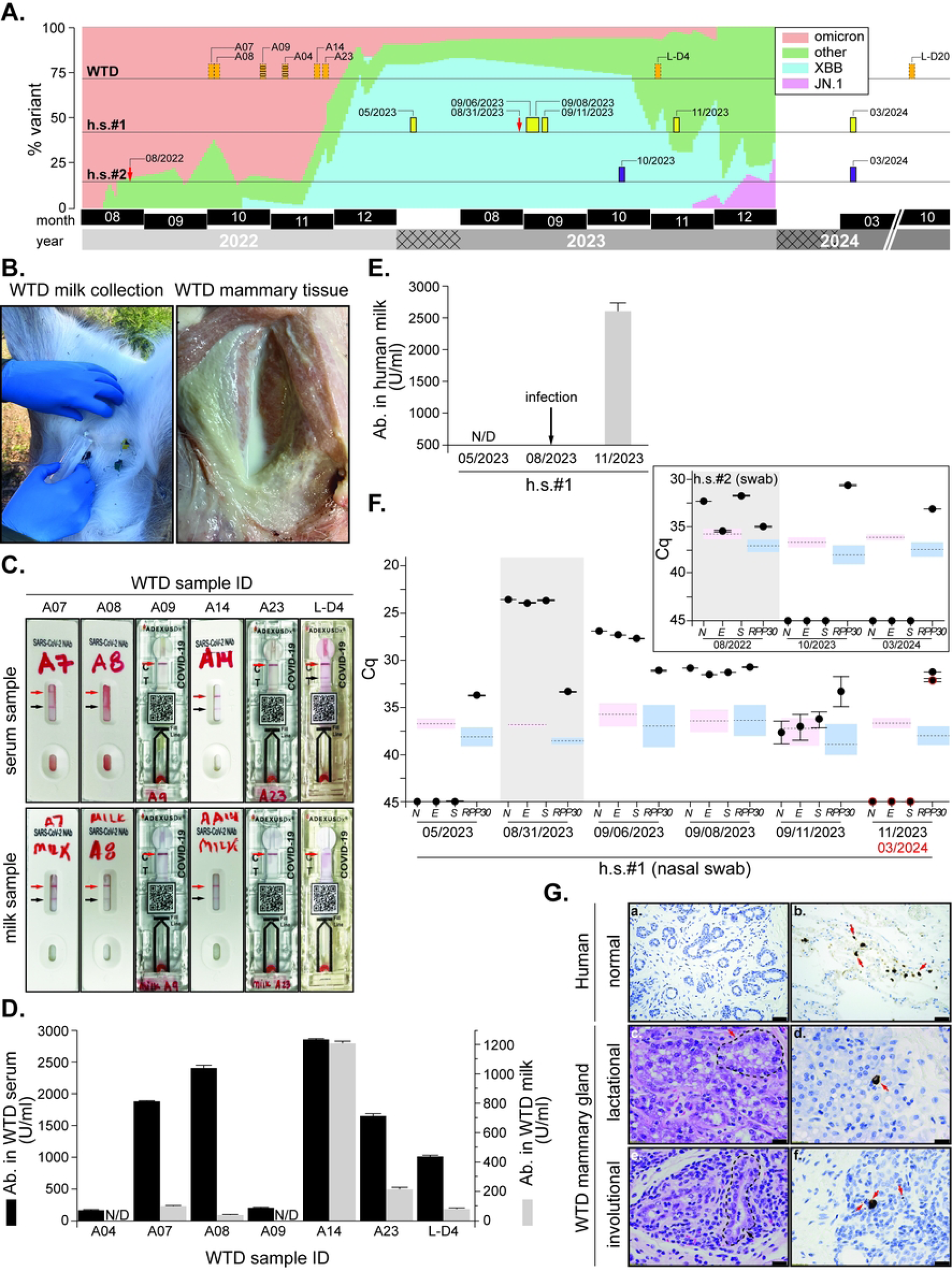
Detection of SARS-CoV-2 antibodies and viral RNA in white-tailed deer (WTD) and human specimens (h.s.). (**A**) Timeline showing SARS-CoV-2 variant circulation in central Virginia (2022-2024) based on whole-genome sequencing of 654 positive specimens. Variants are categorized as Omicron (orange), XBB (blue), JN.1 (pink), and other (green). Horizontal tracks indicate the source of samples WTD (top) and human subjects (middle and bottom labeled h.s.#1 and h.s.#2, respectively). Sample collection timepoints for deer [orange dashed bars, sample IDs (collection date): A04 (11/22), A07 (10/22), A08 (10/22), A09 (10/22), A14 (11/22), A23 (11/22), L-D4 (11/23), and L-D20 (10/24)] and human subjects (h.s.#1 in yellow, h.s.#2 in purple) are indicated relative to variant prevalence in the region. Red arrows indicate times at which the human subject sample yield a positive result by RT-qPCR. The x-axis corresponds to the timeline (month in black and year in shades of gray) from mid-2022 through early 2024 when collection took place. Crossed patterns indicate the time of the year at which collection did not occur. (**B**) Milk collection from a lactating doe (left) and surgically incised mammary gland displaying homogenous tissue with milk expression (right). (**C**) Lateral flow immunochromatography results for SARS-CoV-2 antibodies in serum (upper panel) and milk (lower panel) from six WTD specimens (A07, A08, A09, A14, A23, L-D4). White and clear cassettes are Genscript and ADEXUSDx COVID-19 (from NOWDx, Inc.), respectively. Red arrow indicates a valid positive control test. Black arrow indicates the position of a band if positive for SARS-CoV-2 antibodies. (**D**) Quantitative ELISA results showing SARS-CoV-2 neutralizing antibody levels in deer serum (black bars, left axis) and milk (gray bars, right axis). Values represent mean ± SEM in U/ml. Samples A04 and A09 were below detection limit (N/D, <15 U/ml). (**E**) Antibody levels in human milk from h.s. #1 measured by ELISA at three timepoints: pre-infection (05/2023, N/D), post-infection (08/2023), and late convalescence (11/2023). Arrow indicates timing of documented infection. Error bars indicate standard deviation (SD) among technical replicates for each milk sample. (**F**) RT-qPCR analysis of human nasal swabs showing SARS-CoV-2 RNA detection (*N*, *E*, *S* genes) and human *RPP30* control. Cq values are plotted for human subject #1 at multiple timepoints pre- and post-infection. Inset: RT-qPCR results for human subject #2 at the time of infection. Plotted values indicate mean ±S D of a triplicate. Detection thresholds and corresponding SD are indicated as dashed lines with pink and blue shades for viral and RPP30 genes, respectively. Cq values below the threshold indicate a positive result (note the y-axis is inverted). Gray shading indicates an active infection period. (**G**) Normal mammary gland histological examination (SARS-CoV-2 negative) (a) and post-mortem SARS-CoV-2 nucleocapsid-positive immunohistochemistry in lung (b, arrows). H&E stained mammary gland lobules from a lactational (c) and a non-lactating involutional (e) WTD does. H&E-stained mammary gland lobules from lactating (c) and non-lactating involutional (e) WTD does. Note coalescing alveoli composed of vacuolated secretory cells within the lactating lobule and inactive alveoli with intervening stroma and lymphocytes within the involutional lobule (alveolus - dashed outline). Rare SARS-CoV-2-positive histiocytes (arrows) are present within lactating (d) and non-lactating (f) mammary gland lobules. Hematoxylin and eosin (H&E), original magnification 40x (c, e); SARS-CoV-2 immunohistochemistry original magnification 20x (a, b) and 40x (d, f). Images captured with Canon Eos 70D (a, b) and Olympus DP 72 (c, d, e, f) cameras. Scale bars represent 50 μm.

### Detection of SARS-CoV-2 antibodies in WTD and human samples

Following SARS-CoV-2 infection, humans develop humoral immune responses characterized by initial IgM production followed by IgG and IgA isotypes, with IgA providing crucial mucosal protection [35]. These neutralizing antibodies have been detected in various human fluids, including saliva, respiratory secretions, serum, and breast milk [36–39]. It is speculated that these antibodies reduce the infectivity of human secretions and viral transmission, while also acting as a form of passive immunity [35, 40, 41]. To determine whether similar immune-protective mechanisms exist in wildlife reservoirs, we analyzed antibody and viral RNA presence in milk and serum from harvested female WTDs (Figure 1B-F).

Milk and mammary gland tissue were obtained from legally harvested lactating does (Figure 1B and Sup. Figure 1) in consultation with the Virginia Department of Wildlife Resources using contamination-prevention protocols (see Materials and Methods). Given the potential for species-specific immune responses and the unknown kinetics of antibody production in WTD, we employed multiple detection approaches to comprehensively assess SARS-CoV-2 antibody presence (Figure 1C-D). These included field-deployable lateral flow immunochromatography (GenScript SARS-CoV-2 Neutralizing Antibody Detection kit, ADEXUSDx COVID-19 Test and NOVODIAX CoNAb SARS-CoV-2 Neutralizing Antibody Lateral Flow Test) and laboratory-based ELISA (Abcam COVID-19 S-Protein S1RBD Human IgG ELISA kit). The ADEXUSDx test detects total SARS-CoV-2 antibodies (IgM, IgG, and IgA), while NOVODIAX and GenScript assays specifically identify neutralizing antibodies capable of blocking spike protein-angiotensin-converting enzyme 2 (ACE2) receptor interactions. The ELISA provided quantitative measurement of anti-RBD (receptor binding domain) IgG levels (Figure 1C-E).

Analysis of deer samples revealed variable antibody responses (Figure 1C-D). Three deer (A07, A08, A14) tested positive across all platforms in both serum and milk. Serum antibody concentrations were 691.8 ± 36.5, 2396.8 ± 33.4, and 2854.3 ± 7.9 U/ml, respectively, with corresponding milk concentrations of 136.8 ± 14.1, 74.7 ± 4.5, and 1237.3 ± 147.7 U/ml. Sample L-D4 showed positive lateral flow results only in serum but demonstrated detectable IgG by ELISA in both serum (948.2 ± 13.4 U/ml) and milk (157.6 ± 22.6 U/ml). Sample A23, while negative by lateral flow assay, yielded measurable antibody levels by ELISA (serum: 630.5 ± 47.5 U/ml; milk: 227.5 ± 5.9 U/ml). Two samples (A04, A09) remained negative across all assays.

To establish comparative context for these wildlife findings, we examined antibody dynamics in human milk samples with known infection histories. Human breast milk from h.s.#1 was collected at three timepoints: pre-infection (05/2023), during confirmed infection (08/2023), and post-infection (11/2023 and 03/2024). Post-infection IgG levels reached 2441.1 ± 151 U/ml (11/2023) and 488.2 ± 98 U/ml (03/2024) for h.s.#1, while h.s.#2 showed undetectable milk antibodies one year after infection. Neither subject received vaccine boosters during sample collection (Figure 1E). These findings align with reports that SARS-CoV-2 neutralizing antibodies in breast milk are highly time-dependent, peaking 1-3 months post-infection and waning to undetectable levels within a year without booster exposure [42, 43].

Our results provide the first evidence of SARS-CoV-2–specific antibodies, including neutralizing IgG, are present in the milk of a wildlife species, demonstrating that lactating WTD mount mucosal and systemic immune responses comparable to those observed in humans. The presence of virus-blocking antibodies in milk suggests the potential for passive immunity in fawns through suckling and reveals an unrecognized immune barrier that may influence viral maintenance and transmission within deer populations.

### Viral RNA detection

Next, we tested for the presence of SARS-CoV-2 particles in milk from both WTD and human specimens, as well as from nasal swabs obtained at the indicated sample-collection timepoints (Figure 1A). Nasal swabs collected from h.s.#1 and h.s.#2 at three timepoints, pre-SARS-CoV-2 exposure (05/2023 and 08/2022 for h.s.#1 and h.s.#2, respectively), during exposure (08/31/2023, 09/06/2023, 09/08/2023 for h.s.#1), and post-exposure (11/2023 and 03/2024 for h.s.#1 and h.s.#2, respectively), were analyzed using RT-qPCR targeting the SARS-CoV-2 N, E, and S genes, as previously described [34]. Similarly, nasal swabs from WTDs were obtained at the time of milk collection and analyzed using the same RT-qPCR protocol.

Amplification of SARS-CoV-2 genes were detected in human nasal swab samples but not in milk samples from h.s.#1 collected on 08/31/2023, 09/06/2023, and 09/11/2023, nor in samples from h.s.#2 collected in 08/2022, which correspond to the dates surrounding the first positive diagnostic results for both volunteers. Swabs collected before infection (h.s.#1, 05/2023) or after infection (h.s.#1, 11/2023; h.s.#2, 10/2023 and 03/2024) yielded negative RT-qPCR results (Figure 1F and inset). Viral particles were not detected in nasal swabs or milk samples from WTD using both transcription-mediated amplification (TMA) [Hologic Panther, [44]] and RT-qPCR shortly after collection, indicating that although WTD produced SARS-CoV-2–specific antibodies, there was no evidence of active viral shedding at the time of sampling.

### Histopathological Analysis

Mammary gland tissue examination revealed differences between physiological states (Figure 1G). Immunohistochemistry detected rare SARS-CoV-2-positive histiocytes, likely representing a consequence of incidental histiocytic migration rather than primary mammary gland infection. No evidence of active viral replication was observed in mammary epithelial cells.

The absence of detectable viral particles in WTD milk despite the presence of neutralizing antibodies (Figure 1) raised important questions about the mechanisms underlying viral clearance in this biological matrix. Therefore, to determine whether WTD milk possesses intrinsic antiviral properties that could explain the absence of detectable virus in field samples, and to identify the temporal window during which viral particles might remain stable during acute infection phases, we investigated SARS-CoV-2 stability in WTD *versus* human milk under controlled conditions.

### Temperature-dependent viral stability analysis

For all our *in vitro* studies, we chose to work with three variants of SARS-CoV-2 prevalent in the region and that represent distinct evolutionary stages: PANGO lineage A (original variant), B.1.1.7 (Alpha, characterized by enhanced infectivity due to spike protein mutations), and BA.1.1.529 (Omicron, highly mutated variant with immune evasion capabilities and global dominance) [2, 8, 45]. For comparison, milk samples from WTD A14 and human subject h.s.#1 (pre-infection, 05/2023) were spiked with equal copies of inactivated, variant-specific virus and incubated at room temperature (22°C), physiological temperature (37°C), or pasteurization temperature (63°C) (Figure 2). Aliquots were collected at different times for RT-qPCR analysis (Figure 2).

**Figure 2.**
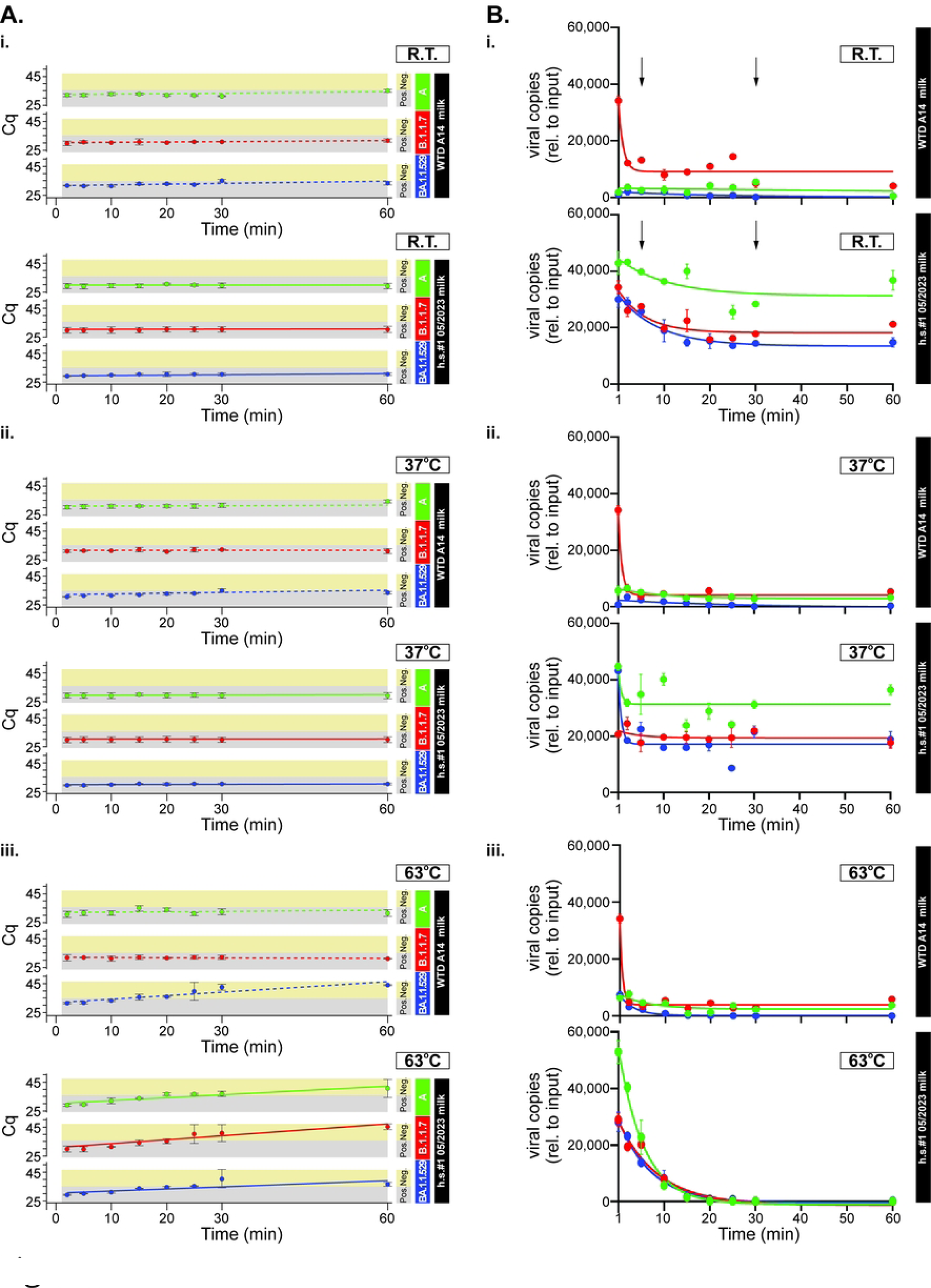
Thermal stability of SARS-CoV-2 variants in human and WTD milk. (**A**) Time-course RT-qPCR analysis of SARS-CoV-2 variant stability in milk at room temperature (RT, 22°C, panels i), 37°C, (physiological temperature, panels ii) and 63°C (pasteurization temperature, panels iii). Human and WTD milk samples were spiked with A (green), B.1.1.7/Alpha (red), or BA.1.1.529/Omicron (blue) variants and incubated for up to 60 min. Cq values indicate remaining viral RNA levels over time, with higher Cq values representing lower viral loads. For each experiment, the cut-off threshold value for the detection of the *N* gene separates regions for which Cq values indicate a positive versus a negative result (gray- and yellow-shaded areas, respectively). Each plotted point is represented as the mean Cq ± SD. A linear relationship has been drawn between the points within each panel to show progressive degradation. Cq cut-off for the N gene was 36.73 ± 0.74 **(B)** Quantitative viral copy analysis showing viral RNA stability expressed as copies relative to input over time. Panels i, ii, and iii correspond to incubation temperatures of RT (22°C), 37°C, and 63°C, respectively. Upper panels show WTD A14 milk; lower panels show human h.s.#1 (05/2023) milk. Viral variants are color-coded as in panel A: A (green), B.1.1.7/Alpha (red), and BA.1.1.529/Omicron (blue). Data points represent mean viral copies ± SD. Note the rapid decline in viral copies in WTD milk compared to stable levels maintained in human milk at physiological temperatures.

Temperature-dependent stability analysis revealed important species-specific differences in viral RNA preservation (Figure 2A-B). In human milk, all three variants demonstrated remarkable stability at room temperature and 37°C, with Cq values remaining consistently below the experimental cut-off for the N gene [defined as in [34]] throughout the 60 min incubation period (Figure 2A.i-ii and 2B.i-ii,). Viral copy numbers in human milk showed initial slight declines but then plateaued relative to input, maintaining stable levels across the experimental timeframe (Figure 2B.i-ii). Conversely, WTD milk exhibited rapid degradation of all three viral variants (A, B.1.1.7, and BA.1.1.529) under identical conditions at both room temperature and 37°C. In WTD milk, viral copy numbers displayed a rapid decline in the first 10 min, dropping from initial levels to near-undetectable, with RNA from all variants falling below detectable levels (Cq >40) within 30-40 min at physiological temperatures (Figure 2B.i-ii)

At pasteurization temperature (63°C), both milk types showed rapid viral inactivation, with all variants becoming undetectable within 20-30 min (Figure 2A.iii and 2B.iii). However, the kinetics of thermal degradation differed between species, with WTD milk showing more rapid initial degradation compared to human milk. These findings demonstrate that WTD milk possesses intrinsic factors that actively degrade viral RNA under physiological conditions, while thermal treatment at pasteurization temperatures effectively inactivates viral particles in both species’ milk, albeit through different kinetic patterns.

### Viral RNA stability patterns reflect intrinsic milk properties

Short-term viral stability assessment confirmed that rapid RNA degradation in WTD milk is a consistent phenomenon across different collection timepoints and individual animals. Both WTD samples A23 and L-D4, collected approximately one year apart, demonstrated similar degradation patterns for all three variants (Figure 3A). Similarly, human milk from h.s.#1 showed consistent viral preservation properties in both pre-infection (05/2023) and post-infection (11/2023) samples, despite physiological changes associated with SARS-CoV-2 infection and lactation status. This temporal consistency indicates that the differential viral stability reflects intrinsic species-specific milk properties rather than individual variation or infection-related changes.

**Figure 3.**
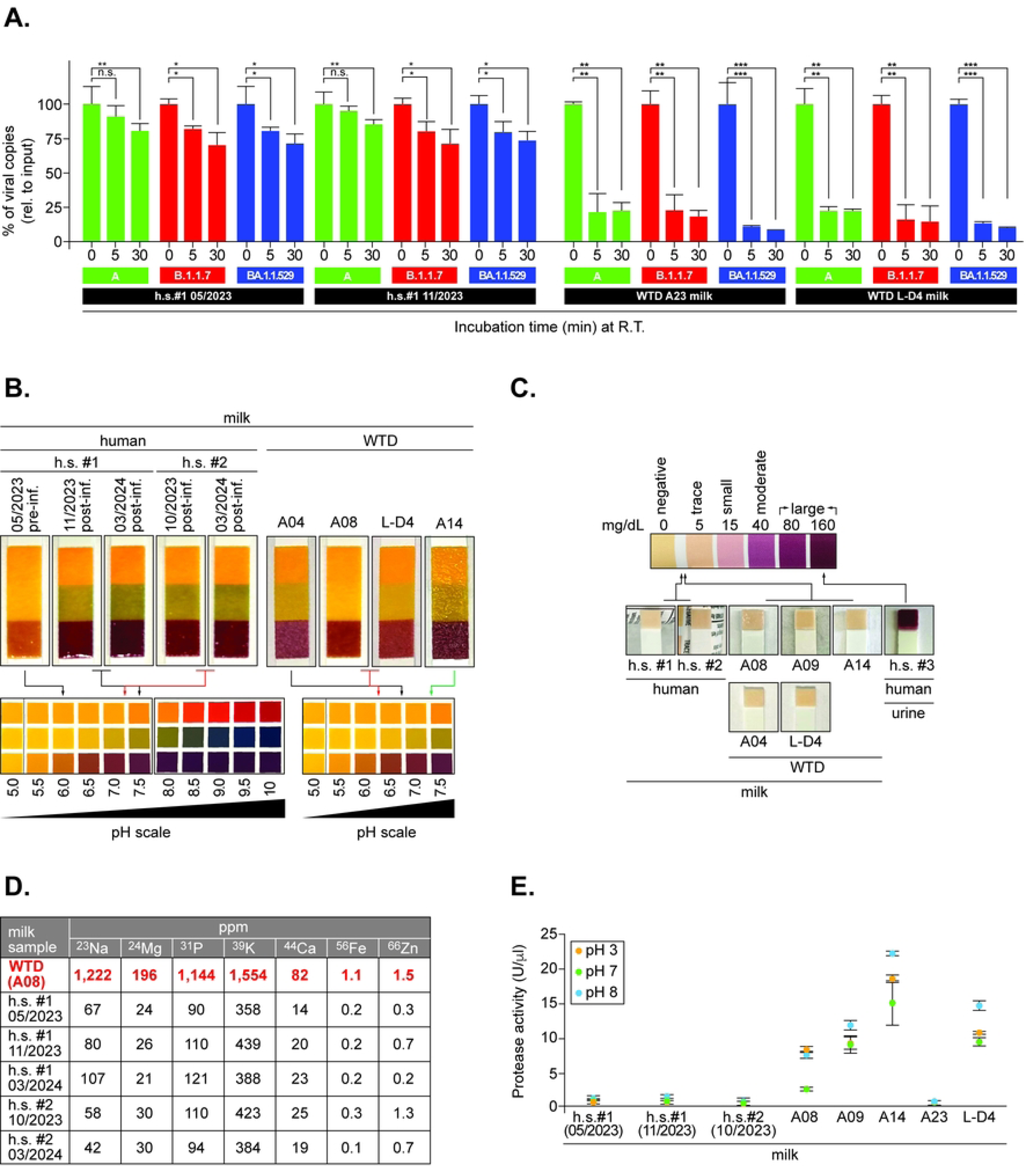
Viral RNA degradation in milk and effects of protease inhibition. (**A**) Comparative viral RNA stability in human and WTD milk. Samples were spiked with A (green), B.1.1.7 (red), or BA.1.1.529/Omicron (blue) variants and incubated at room temperature for 5- or 30-min. Results show percent viral RNA recovery relative to input. Human milk (h.s.#1 05/2023, h.s.#1 11/2023) maintained higher RNA stability compared to WTD samples (A23, L-D4). (**B**) pH determination of human and WTD milk samples using colorimetric indicator strips. Color changes correspond to pH values ranging from 5.0-10 as indicated by the color scale. Human milk samples (h.s.#1, h.s.#2) and WTD samples (A04, A08, L-D4, A14) are shown. Dates indicate the time at which the corresponding milk sample was collected based on the information displayed in Figure 1A. For each milk sample, the pH test strip (upper strip) is shown alongside the manufacturer’s reference color scale (below) to allow visual comparison. The strip color for each sample was matched to the reference scale, and the pH was recorded to the nearest 0.5 unit based on the best color match. (**C**) Ketone body detection in milk samples using PrimeScreen test strips. Color intensity correlates with ketone concentration (from 0 to >160 mg/dl scale). Human milk (h.s.#1, h.s.#2) and WTD milk (A08, A09, A04, L-D4, A14) are shown. A human urine sample (h.s.#3) from a known ketone-positive donor is included as a positive control. (**D**) Elemental composition analysis by inductively coupled plasma mass spectrometry (ICP-MS) showing concentrations (ppm) of seven key minerals (Na, Mg, P, K, Ca, Fe, Zn) in human milk samples (h.s.#1 at multiple timepoints, h.s.#2) and WTD milk (A08). Values in red indicate ≥5-fold higher concentrations in WTD milk compared to human milk. (**E**) Protease activity measurements in human (h.s.#1, h.s.#2) and WTD milk samples (A08, A09, A14, A23, and L-D4) at pH 3, 7, and 8 using fluorometric assay. Activity values were normalized and reported as corrected fluorescence units (U/μl). Error bars represent mean ± SD from triplicate measurements.

### Biochemical environment characterization: pH and metabolic profiling in milk

We hypothesized that species-specific differences in milk pH could contribute to the differential viral stability patterns observed between human and WTD milk. pH measurements using colorimetric strips revealed species-specific differences and temporal changes reflecting various physiological conditions (Figure 3B). A pH 6.0 was measured in human milk from h.s.#1 pre-infection (05/2023, 11 months postpartum), consistent with mature human milk profiles [46], which increased to approximately 7.25 post-infection (11/2023), possibly reflecting immune-mediated changes or infection-related mammary gland alterations. Notably, viral RNA stability was maintained in human milk despite the slightly acidic pre-infection pH (Figure 3B), indicating that pH alone does not account for the preservation observed in human samples. WTD milk samples displayed neutral to slightly alkaline pH values, creating a different chemical environment that may influence enzymatic activity and viral stability (Figure 3B).

We further hypothesized that altered metabolic states, as indicated by ketone body presence, could reflect physiological stress or metabolic changes that might influence milk composition and antiviral properties. Ketone body analysis using test strips revealed minimal levels in both human and WTD milk samples compared to a positive control (human urine from subject h.s.#3) (Figure 3C). The consistently low ketone concentrations across all milk samples indicate normal metabolic profiles in both species, ruling out ketosis-related factors as contributors to the observed differences in viral stability.

### The composition of mineral elements substantially varies among species

To identify species-specific differences in mineral composition that may influence SARS-CoV-2 RNA stability, we performed inductively coupled plasma mass spectrometry (ICP-MS) on milk samples from both species (Figure 3D). Analysis of multiple human milk samples collected at different timepoints and lactation stages, including h.s.#1 samples at 05/2023 (pre-infection), 11/2023 (post-infection), and 03/2024, as well as h.s.#2 samples at 10/2023 and 03/2024, revealed remarkably consistent mineral profiles despite temporal and physiological variations (Figure 3D). In contrast, WTD milk (sample A08) demonstrated markedly elevated concentrations across seven key elements. Sodium, magnesium, phosphorus, and potassium showed the most striking differences, with concentrations of 1,222, 196, 1,144, and 1,554 ppm respectively, representing 5-20-fold elevations compared to human milk samples. Iron, calcium, and zinc were similarly elevated in deer milk. The consistency of low mineral concentrations across all human samples, regardless of collection timepoint or infection status, highlights these differences as intrinsic species characteristics rather than individual or temporal variations. These elevated ionic concentrations in WTD milk create a distinct biochemical environment that may contribute to enhanced enzymatic activity and altered viral stability.

### Species-specific protease activity and pH dependencies

To assess species- and pH-dependent differences in proteolytic activity, we measured total protease activity in various WTD and human milk using the Amplite Protease fluorometric assay at pH 3, 7, and 8 (Figure 3E). Human milk samples demonstrated notable consistency both between individuals and across infection status. Subject h.s.#1 showed minimal protease activity pre-infection (05/2023), with slight increases post-infection (11/2023), particularly at pH 7. Subject h.s.#2 (10/2023) exhibited comparable activity levels compared to h.s.#1 samples. All human milk samples maintained relatively low proteolytic activity across the pH range tested, with optimal activity consistently observed at physiological pH 7 (Figure 3E). By contrast, WTD milk samples (A08, A09, A14, L-D4) exhibited substantially higher protease activity than any human sample across all pH conditions tested, with activity levels 3-5 fold greater than human milk (Figure 3E). WTD samples showed consistent pH-dependent patterns with peak activity at pH 7, indicative of alkaline-active proteases (e.g., plasmin-plasminogen system and matrix metalloproteinases). Notably, sample A23 demonstrated protease activity levels more comparable to human samples (Figure 3E), yet achieved complete viral RNA degradation similar to other WTD samples (Figure 3A). This observation suggests that proteolytic activity alone may not fully account for the antiviral properties of WTD milk, and that the elevated ionic concentrations (Figure 3D) and altered pH environment (Figure 3B) may provide suitable conditions for viral RNA degradation even when protease levels are reduced. The synergistic effect of multiple biochemical factors, including ionic strength, pH, and proteolytic activity, likely creates a threshold environment where even moderate increases in any single factor can achieve effective viral inactivation. This finding highlights the multifactorial nature of the antiviral environment in WTD milk and suggests that complete viral degradation can be achieved through different combinations of these biochemical mechanisms.

### Mineral supplementation effects on SARS-CoV-2 stability

We hypothesized that the elevated mineral concentrations detected in WTD milk could be responsible, at least in part, for the rapid viral instability observed in these samples. To test this hypothesis, we reasoned that supplementing human milk (which has lower mineral concentrations) with individual minerals or mineral cocktails at levels equivalent to those found in WTD milk would provide mechanistic insights into ionic-mediated viral degradation. Accordingly, we supplemented human milk (h.s.#1, pre-infection 05/2023) with individual minerals at concentrations equivalent to those detected in WTD milk (Figure 4A). Each mineral was tested individually across three temperature conditions (room temperature, 37°C, 63°C) with all three viral variants (A, B.1.1.7, BA.1.1.529) to assess mineral-specific and temperature-dependent effects on viral stability (Figure 4A). At room temperature (Figure 4A, left panels), individual mineral supplementation revealed variant-specific susceptibilities. The B.1.1.7 variant (red) demonstrated significant sensitivity to mineral addition, showing substantial reductions in viral RNA recovery with all tested minerals compared to control conditions. In contrast, both the A variant (green) and BA.1.1.529 (blue) maintained viral copy numbers comparable to input values across mineral supplementation conditions, indicating greater stability under these conditions. At 37°C and 63°C, the differential mineral sensitivity became more pronounced, with B.1.1.7 showing the greatest susceptibility to mineral-induced degradation across all temperature conditions, while A and BA.1.1.529 maintained relatively better stability compared to B1.1.7 at similar temperature (Figure 4A, middle and right panels). The relative stability of A and BA.1.1.529 variants under identical ionic conditions suggests that their respective mutation profiles confer either enhanced structural resilience or different ion-binding characteristics that do not compromise viral particle integrity under the tested conditions.

**Figure 4.**
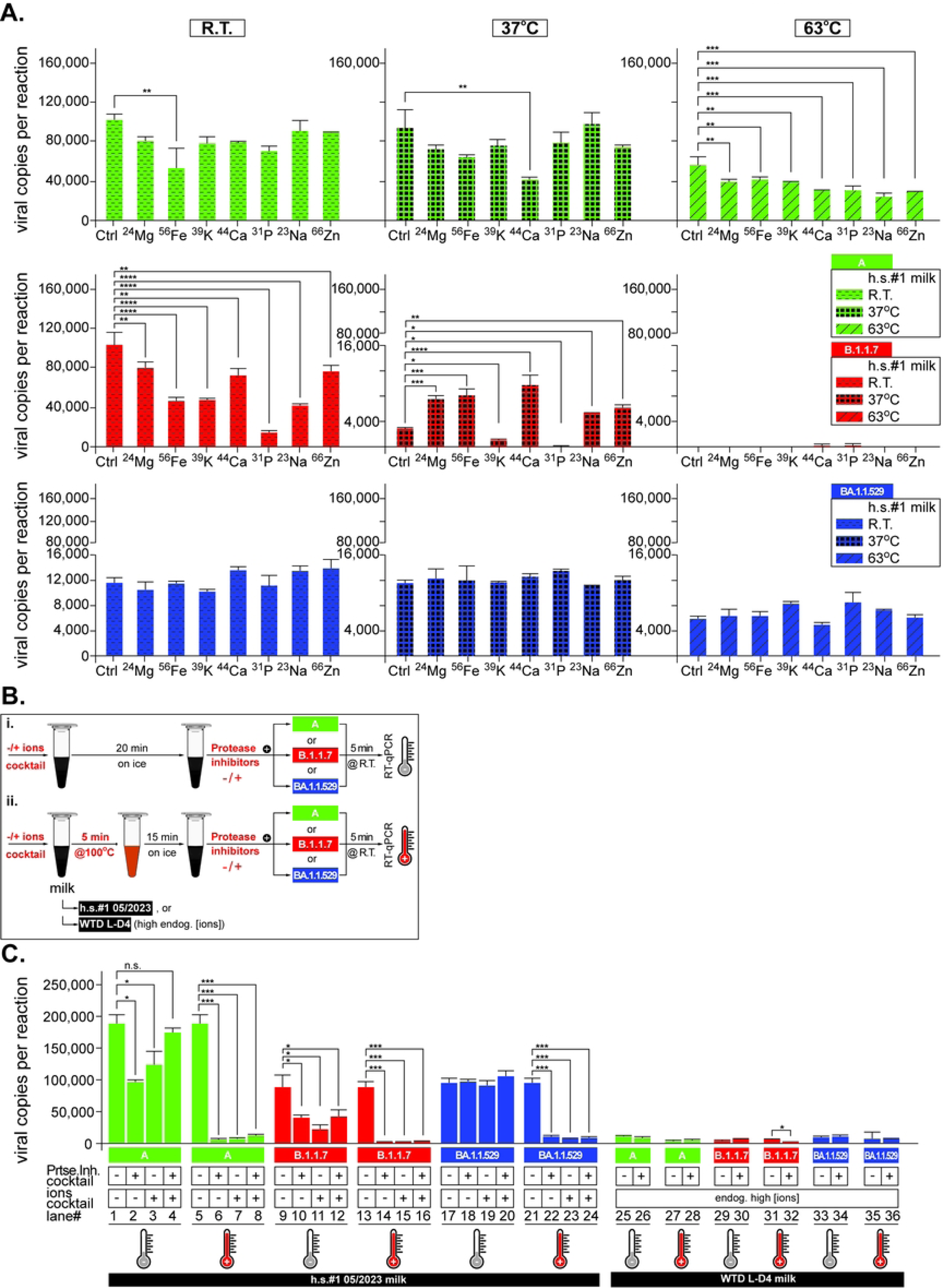
Mineral supplementation influences SARS-CoV-2 variant stability in milk. (**A**) Human milk (h.s.#1 05/2023) was supplemented with individual ions (²³Na, ²⁴Mg, ³¹P, ³⁹K, ⁴⁴Ca, ⁵⁶Fe, ^6^⁶Zn) at concentrations equivalent to those found in WTD milk as described in the Materials and Methods section. Samples were spiked with A (green), B.1.1.7/Alpha (red), or BA.1.1.529/Omicron (blue) variants and incubated at room temperature (RT, 22°C), 37°C, or 63°C. Viral copy recovery is shown as copies per reaction (mean ± SD). Control (Ctrl) represents unsupplemented human milk. (**B**) Step-by-step experimental workflow diagram for assessing viral RNA degradation in milk under various conditions (e.g., ion supplementation, protease inhibitors). Panel (i) Human milk (h.s.#1) was supplemented, or not, with ion cocktails to mimic the levels found in WTD milk, maintained for 20 min on ice, and then treated, or not, with protease inhibitors before spiking with the indicated virus and performing RT-qPCR analysis as described in the Materials and Methods section. For WTD milk (L-D4), endogenous ion levels exist and only the effect of protease inhibitor addition was monitored. In panel (ii), the experimental protocol includes an intermediate step where samples are incubated at 100°C for 5 min followed by 15 min on ice before adding, or not, protease inhibitors and proceeding as in (i). (**C**) Effects of protease inhibition and ion supplementation on viral RNA degradation. Human milk (h.s.#1 05/2023) and WTD milk (L-D4) were treated with combinations of protease inhibitor cocktail and ion mixture, then spiked with viral variants as indicated in (B). Bar graphs show viral copies per reaction under different conditions (mean ± SD). Temperature symbols indicate room temperature (gray thermometer) or heat treatment at 100°C (red thermometer). In all cases statistical significance is indicated as: *p<0.05, **p<0.01, ***p<0.001, ****p<0.0001.

### Mechanistic analysis of ionic and proteolytic factors contribution to viral stability

Next, we aimed to dissect the relative contributions of the ionic environment and proteolytic activity to viral degradation in milk from both species (Figure 4B-C). Accordingly, we designed controlled *in vitro* experiments testing ion supplementation and protease inhibition in milk samples spiked with each viral variant (Figure 4B, scenario i). The ion cocktail contained all minerals at concentrations equivalent to those found in WTD milk when added to human milk, whereas WTD milk from sample L-D4 relied on its endogenous high ion concentrations.

Results show that the addition of protease inhibitors alone to human milk prior to spiking viral particles does not prevent the A and B.1.1.7 variants from being degraded (Figure 4C, lanes 1 vs 2 and 9 vs 10) but contributed to BA.1.1.529 stability in human milk (Figure 4C, lanes 17 vs 18). Ion cocktail addition primarily impacted the B.1.1.7 variant stability, albeit degradation of A variant is noted (Figure 4C, lanes 1 vs 3), likely through activation of endogenous protease activity, as subsequent protease inhibitor addition partially restored stability (Figure 4C, lanes 1 vs 4), indicating that both ionic stress and proteolytic activity contribute synergistically to viral degradation.

Of note, whereas B.1.1.7 stability remains sensitive to ion addition at room temperature (lanes 9 vs 11), BA.1.1.529 remained largely stable at room temperature within the tested timeframe (Figure 4C, lanes 9 17-20), with minimal impact from ion mixture addition and protease inhibition. This contrasts with the B.1.1.7 sensitivity observed with individual ion supplementation (Figure 4A), suggesting that the combined ionic environment may saturate degradative pathways differently than individual ion stress, or that competitive binding between multiple ions reduces the specific destabilizing effects observed with single mineral supplementation.

In WTD milk sample L-D4, protease inhibitors provided modest protection, but substantial degradation persisted (Figure 4C lanes 25-26, 29-30, and 33-34), confirming that elevated endogenous ionic concentrations alone can promote viral instability independent of proteolytic activity. This finding aligns with studies demonstrating that divalent cations can modulate and, at sufficiently high concentrations, destabilize RNA-protein complexes, and that specific binding of divalent cations on viral surface proteins can alter virion stability and conformational transitions [47–49].

Next, we reasoned that a heat shock treatment (100°C for 5 min) followed by protease inhibitor addition prior to virus spiking would reveal the thermal lability of degradative enzymes. Surprisingly, heat pretreatment substantially contributed to viral degradation across all variants and experimental scenarios (Figure 4C, lanes 5-8, 13-16, and 21-24 for human milk and lanes 27-28, 31-32, and 35-36 for WTD milk) indicating that heat-induced protein denaturation and structural changes in the milk matrix create additional degradative conditions beyond enzymatic activity. This unexpected finding suggests that specific thermal treatment can transiently activate or expose enzymatic activities in milk, and may also promote aggregation-associated disruption of viral particles, consistent with studies showing that heat-dependent activation of zymogen-enzyme systems in milk and heat-induced-induced destabilization of virus particles [50–52]. Our results demonstrate that rapid viral RNA elimination in milk, particularly in wildlife samples, involves complex multifactorial mechanisms that extend beyond simple ionic or proteolytic effects.

### Enzymatic profiling and antimicrobial factor analysis

To characterize the enzymatic diversity contributing to differential antiviral properties, we performed an API-ZYM assay on human and WTD milk (without viral spiking) before and after heat treatment. Human milk exhibited a broad spectrum of enzyme activities, with strong reactions observed for esterases, alkaline and acid phosphatases, lipases (C14), leucine arylamidase, naphthol-AS-BI-phosphohydrolase, β-glucuronidase, N-acetyl-β-glucosaminidase, and α-fucosidase (Figure 5A, top panel). Heat treatment (100°C for 5 min) dramatically reduced enzymatic activities in human milk, with only esterase and phosphohydrolase retaining measurable activity (Figure 5A, second top panel). WTD milk demonstrated a distinct enzymatic profile with different activity patterns that showed variable thermal sensitivity compared to human milk. Specifically, WTD milk exhibited strong baseline activity for alkaline phosphatase, multiple esterases (C4, C8), lipase (C14), leucine arylamidase, and several glycosidases including β-glucuronidase and N-acetyl-β-glucosaminidase (Figure 5A, third panel). However, following heat treatment (100°C for 5 min), WTD milk showed complete loss of all enzymatic activities, demonstrating that all tested enzymes in deer milk were thermolabile and sensitive to heat inactivation (Figure 5A, lower panel). This contrasts with human milk, which retained activity in three enzymes after identical heat treatment. The complete thermal inactivation of all enzymatic activities in WTD milk indicates that the continued viral degradation observed in heated deer milk samples (Figure 4C) could be attributed to non-enzymatic factors, particularly the elevated ionic concentrations and altered chemical environment, as well as potentially heat-activated antimicrobial mechanisms that are triggered at high temperatures, rather than residual enzymatic activity.

**Figure 5.**
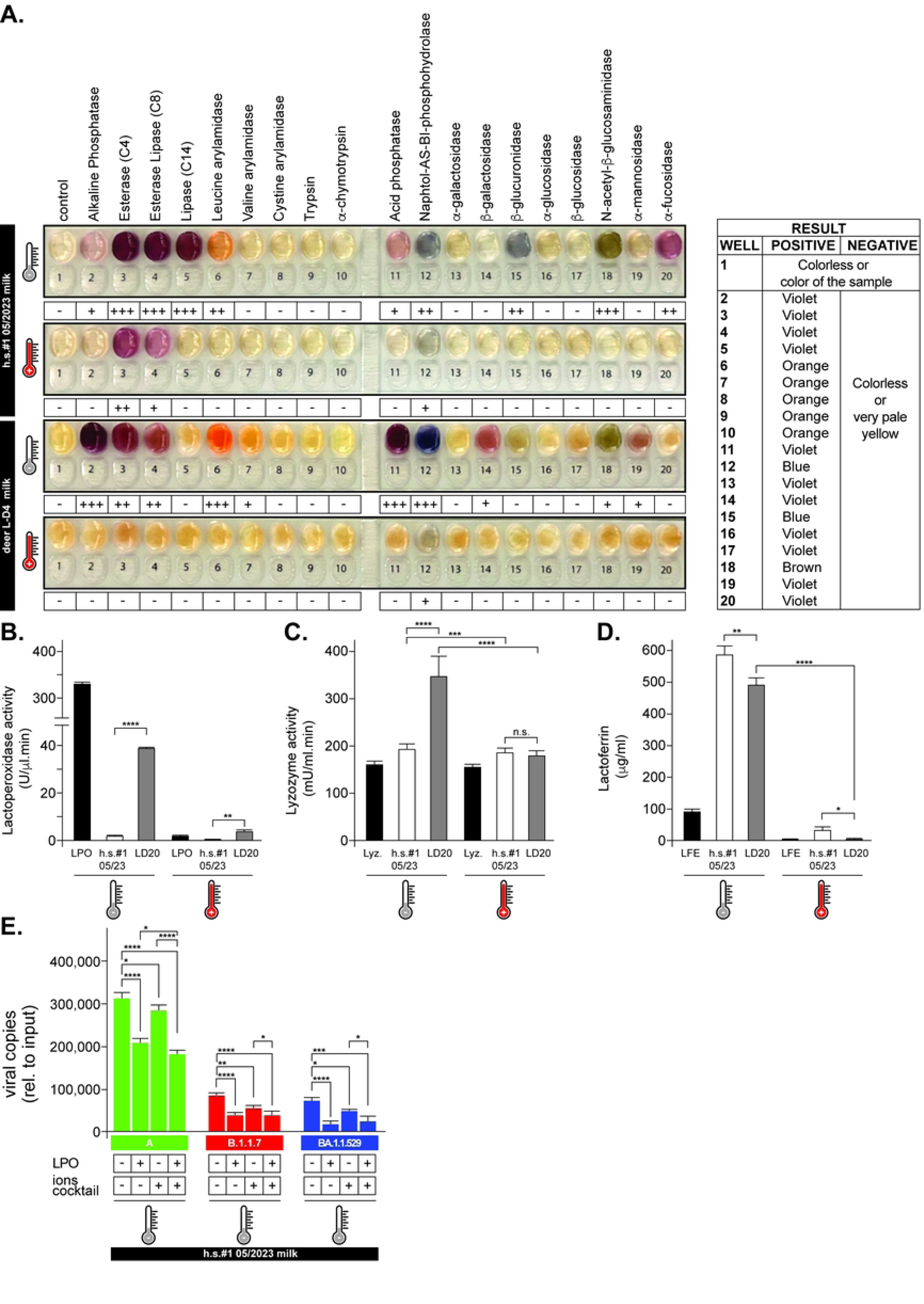
Enzymatic activity profiling and antiviral component characterization in milk. (**A**) API-ZYM enzymatic activity strips showing 20 different enzyme activities in human milk (h.s.#1 05/2023) and WTD milk (L-D4) at room temperature (gray thermometer) and after heating to 100°C (red thermometer). Color intensity indicates enzyme activity levels: negative (-), weak (+), moderate (++), or strong (+++). Enzymes detected include alkaline phosphatase, esterases, lipases, aminopeptidases, glycosidases, and others as labeled. (**B**) Lactoperoxidase (LPO) activity was measured in human milk (h.s.#1 05/2023), WTD milk (L-D20), and commercial LPO standard before and after heat treatment (100°C). Activity is expressed as U/μl·min with error bars showing mean ± SD. (**C**) Lysozyme activity in the same samples and a commercial lysozyme control was measured using fluorometric assay. Activity is expressed as mU/ml·min before and after heat treatment. (**D**) Lactoferrin concentration was measured by ELISA in human and WTD milk samples and commercial lactoferrin standard before and after heat treatment. Concentrations are expressed as μg/mL with error bars showing mean ± SD. (**E**) Functional validation of LPO antiviral activity. Human milk (h.s.#1 05/2023) was supplemented with LPO and/or ion cocktail, then spiked with A (green), B.1.1.7/Alpha (red), or BA.1.1.529/Omicron (blue) variants. Bar graphs show mean ± SD of viral RNA copies recovery relative to input.

To evaluate whether heat treatment affects innate antimicrobial components that contribute to viral degradation, we quantified endogenous lysozyme activity, lactoperoxidase activity, and lactoferrin concentration in untreated and heat-treated milk samples (without viral spiking) from human subject h.s.#1 and WTD sample LD20.

WTD milk demonstrated substantially higher baseline lactoperoxidase activity (∼40 U/μL·min) compared to human milk (<0.01 U/μL·min) (Figure 5B). Heat treatment significantly reduced activity in both milk types; however, the residual activity in WTD milk remained significantly higher than in human milk (Figure 5B). This pattern indicates that while lactoperoxidase is more abundant in WTD milk, it remains thermolabile, suggesting that the enhanced antiviral activity observed in heated deer milk samples must derive from heat-stable factors or heat-activated mechanisms in addition to residual lactoperoxidase function. Next, we quantified lysozyme, a comparatively thermostable antimicrobial enzyme whose resistance to heat inactivation depends strongly on pH and the specific milk matrix [53]. Human milk lysozyme is generally more sensitive to standard pasteurization protocols than lysozyme activity in bovine and other ruminant milks [54]. At room temperature, WTD milk exhibited higher lysozyme activity (345 mU/mL·min) compared to human milk (193 mU/mL·min) (Figure 5C). Heat treatment decreased lysozyme activity to comparable levels for both WTD and human milk and thus, the increased viral instability in WTD milk could not be attributed to the sole activity of lysozyme.

Lastly, we measured lactoferrin activity since this protein exhibits antiviral effects through iron chelation, viral binding, and RNA degradation, and its function is strongly influenced by heat sensitivity and divalent cations [55, 56]. Results show that both species exhibit high baseline levels of lactoferrin, with human milk containing ∼590 μg/mL and WTD milk ∼490 μg/mL (Figure 5D); however, heat treatment caused a dramatic reduction in its activity in both species.

Given the elevated ionic concentrations in WTD milk and their potential role in lactoperoxidase system activation, we tested whether lactoperoxidase supplementation combined with ion cocktail addition could mimic the WTD milk’s antiviral properties in human milk (Figure 5E). To specifically assess synergistic interactions between ions and lactoperoxidase activity, we supplemented human milk with a suboptimal concentration of lactoperoxidase (0.4 U/μL, approximately 10-fold lower than endogenous WTD levels) to determine whether ionic enhancement could amplify the antiviral effect beyond what either component could achieve individually. Results demonstrated that lactoperoxidase addition alone provided moderate antiviral effects, while ion cocktail supplementation enhanced viral RNA degradation. The combination of lactoperoxidase and ions showed synergistic antiviral activity across all variants, mimicking to an extent the degradation pattern observed in native WTD milk (Figure 5E), confirming that ionic concentrations can potentiate lactoperoxidase-mediated viral inactivation.

## Discussion

Our study provides evidence that WTD milk exhibits multifactorial antiviral mechanisms that rapidly inactivate SARS-CoV-2 through synergistic biochemical pathways distinct from those that operate in human milk. The concurrent presence of neutralizing antibodies and intrinsic antiviral properties, mediated by elevated mineral concentrations, enhanced proteolytic activity, and species-specific antimicrobial factors, suggests that milk-borne transmission is unlikely to contribute significantly to viral maintenance in WTD populations. The combination of neutralizing antibodies in milk, viral infection limited to inflammatory histiocytes rather than epithelial cells, and the rapid instability of free viral particles in the WTD milk matrix indicates that lactation represents a pathway of pathogen clearance rather than transmission in this wildlife reservoir species.

Our studies show that in WTD, RT-qPCR of milk targeting *N*, *E*, and *S* genes found no detectable SARS-CoV-2 RNA, supporting prior human data demonstrating viral absence in milk even with positive nasal swabs [17, 57]. Histopathological examination of WTD mammary glands showed no evidence of mammary epithelial infection, with only rare SARS-CoV-2-positive histiocytes detected by immunohistochemistry, consistent with passive histiocytic transport rather than local viral replication (Figure 1). This pattern contrasts markedly with other viral infections where milk-borne transmission is well-documented. For instance, HIV-1 transmission through human breast milk is facilitated by activated, mucosa-homing CD4+ T cells that support viral replication, creating a stable cell-associated viral reservoir that persists even under antiretroviral treatment [58, 59]. Consequently, breastfeeding accounts for more than half of pediatric HIV infections globally in settings without effective prophylaxis [58]. Similarly, Rift Valley fever virus demonstrates prolonged stability and infectivity in ruminant milk, with documented transmission through consumption of contaminated raw milk from infected livestock [28–30]. These contrasting examples highlight the pathogen-specific nature of milk-based transmission dynamics and underscore that SARS-CoV-2 exhibits distinct behavior in mammary environments compared to other viruses with established milk-borne transmission routes.

Variant-specific susceptibility patterns elucidated through mineral supplementation studies provide mechanistic insights into ionic-mediated viral destabilization (Figure 3 and 4). The pronounced sensitivity of B.1.1.7 to mineral stress, contrasted with the relative resilience of the A and BA.1.1.529 variants, indicates that specific mutations modulate viral surface electrostatics in ways that create differential vulnerabilities to ionic perturbation. This differential sensitivity can be explained by structural differences between variants. Computational analyses demonstrate that Omicron variants carry markedly more positive electrostatic potential in the receptor-binding domain than ancestral strains (e.g. A), driven by multiple charge-increasing mutations (Q493R, Q498R, T478K, and Y505H) that add cationic residues near the receptor-binding motif [60, 61]. The B.1.1.7 variant, while exhibiting moderate increases in positive charge and furin site modifications, displays a smaller overall electrostatic shift compared to Omicron’s extensive remodeling [62, 63]. These distinct charge patterns likely generate variant-specific responses to ionic environments, as both theoretical and experimental studies demonstrate that ionic strength strongly modulates electrostatic interactions and virus–surface behavior by altering zeta potential and attachment energies as demonstrated for MS2, Qβ, and Norwalk viruses [64, 65]. Such electrostatic modifications can influence affinity for the ACE2 receptor and lipid rafts, fusogenic potential, and pathogenicity, while also contributing to increased immune escape [65]. Similar principles apply to other viral systems: among adeno-associated viruses (AAVs), pH and ionic strength strongly influence electrostatic interactions and induce conformational shifts that affect capsid–capsid interactions and viral aggregation [66], findings that are particularly relevant given that AAVs are widely used as viral vectors in gene therapy, where formulation stability is critical for commercial therapeutic success [67]. The 5-20-fold elevation in essential minerals found in WTD milk might create an ionic milieu that destabilizes the SARS-CoV-2 viral particles through direct electrostatic disruption of viral envelope proteins and ribonucleoprotein complexes, independent of enzymatic degradation pathways. The observed differential sensitivity suggests that B.1.1.7 variant’s moderate electrostatic modifications may create vulnerabilities to divalent cation interference, while Omicron’s extensive positive charge redistribution may inadvertently confer stability under hyperosmotic conditions through enhanced electrostatic buffering capacity.

The multifactorial nature of the viral degradation in WTD milk, evidenced by persistent antiviral activity despite selective inhibition of individual mechanisms, indicates evolutionary redundancy ensuring pathogen inactivation across diverse physiological states. The observation that specimen A23 achieved complete viral elimination despite exhibiting protease activity levels comparable to human milk demonstrates that no singular mechanism accounts for the antiviral phenotype (Figure 3). Rather, synergistic interactions between ionic stress, pH modulation, proteolytic activity, and antimicrobial effectors create a threshold environment wherein multiple pathways provide compensatory antiviral capacity.

The paradoxical enhancement of viral RNA degradation following thermal treatment suggests activation of cryptic antimicrobial mechanisms or exposure of latent ribonuclease activities within the milk matrix (Figs. 2 and 4). This phenomenon has significant implications for processing safety protocols, as conventional thermal inactivation methodologies developed for bovine milk may inadequately predict viral behavior in cervid milk matrices. The complete thermal ablation of detectable enzymatic activities coupled with sustained viral degradation indicates substantial contributions from heat-activated non-enzymatic mechanisms.

From a public health perspective, the identification of neutralizing antibodies and intrinsic antiviral mechanisms in cervid milk suggests that the risk of milk-borne zoonotic transmission may be lower than previously assumed; however, observed variant-specific differences in ionic susceptibility underscore the need for strain-specific risk assessment, as emergent variants may exhibit altered stability within cervid milk matrices. These findings have important implications for the commercialization of deer-derived products, particularly milk, which is an exceptionally nutrient-dense secretion with high concentrations of fat (approximately 10-12%), protein (7-8%), and total solids (24-27%), often exceeding those of cow, sheep, and goat milk, and is enriched in calcium, phosphorus, zinc, branched-chain fatty acids, and α-linolenic acid [21, 68]. Its elevated buffering capacity and the presence of bioactive whey proteins, including lactoferrin, whose hydrolysates exhibit strong antibacterial activity against foodborne pathogens, further support its potential value in functional foods and nutraceutical applications [23]. Consequently, the safe and effective commercialization of deer milk will require processing strategies tailored to its distinctive biochemical composition to ensure robust pathogen inactivation while preserving endogenous antimicrobial activity.

Ecologically, the presence of neutralizing antibodies in WTD milk provides evidence for passive immunization in wildlife populations, with the potential to modulate population-level susceptibility patterns and viral persistence dynamics. The temporal stability of both humoral responses and antiviral activity across collection intervals and among individuals suggests that these features represent intrinsic species-level characteristics rather than stochastic variation. This consistency has important implications for future epidemiological studies, as maternal antibody transfer may generate age-structured immunity patterns that influence population susceptibility and viral maintenance [e.g., in human milk [69–71]]. Accordingly, our findings suggest that wildlife milk may function as an active immunological interface with potential implications for pathogen transmission dynamics, where the observed species-specific antiviral mechanisms likely reflect evolutionary adaptations to pathogen exposure. While our insights into WTD milk biochemistry provide a foundation for understanding milk-based immunity in wildlife, broader investigations across diverse species and larger sample sizes will be necessary to fully elucidate the role of lactational immunity in wildlife reservoir dynamics and zoonotic risk management at wildlife-human interfaces.

## Materials and Methods

### Contamination-prevention protocols for WTD milk collection

To ensure sample integrity and to prevent cross-contamination during field collection of white-tailed deer specimens, research personnel implemented comprehensive biosafety protocols adapted for outdoor conditions. All field collection activities were conducted using personal protective equipment including disposable nitrile gloves, face masks, and protective clothing that were changed between each animal specimen. Collection instruments (syringes, collection tubes, scalpels) were single-use and sterile, with separate sets designated for each sample type (milk, serum, tissue). Milk collection surfaces and teats were cleaned with 70% isopropanol prior to sampling, and collection was performed using aseptic technique to minimize environmental contamination. All samples were immediately placed in sterile, labeled containers and maintained on ice in portable coolers during field transport to prevent degradation and bacterial growth. Field collection areas were sanitized between animals using 10% bleach solution, and personnel underwent hand sanitization protocols between specimen collections. To eliminate the possibility of contamination from research personnel, all team members involved in sample collection provided nasal swab samples for SARS-CoV-2 testing by antigen lateral flow assay or RT-qPCR analysis during the WTD hunting seasons. Sample processing and storage followed standard laboratory biosafety procedures upon return from the field, with samples handled in designated areas to prevent cross-contamination and stored at appropriate temperatures (-20°C for short-term, -80°C for long-term storage) until analysis. Chain of custody documentation was maintained throughout the field collection and laboratory storage process to ensure sample traceability and integrity.

### Milk collection and processing

Milk samples were obtained from WTD does during the 2022-2024 hunting seasons in consultation with the Virginia Department of Wildlife Resources. Samples were obtained by sequentially amputating 5.0 mm of the distal teat aspect and manually expressing milk into sterile conical tubes. Total milk volumes were aliquoted and stored at -20°C and -80°C until analysis. Mammary gland tissue was excised from all lactating, as well as representative non-lactating, does and fixed in 10% neutral buffered formalin for histological evaluation. Human milk samples were collected from consenting lactating volunteers (h.s.#1 and h.s.#2) at specified timepoints (for h.s.#1 05/2023 was pre-infection and 11 months after delivery) and stored similarly under Virginia Tech IRB (IRB#24-474) approval.

### Histopathological analysis

The WTD mammary gland tissues were fixed in 10% neutral buffered formalin, paraffin-embedded, sectioned at 5 μm thickness, and stained with hematoxylin and eosin using the Sakura TissueTek Prisma system. Immunohistochemistry was performed using mouse monoclonal antibody against SARS-CoV/SARS-CoV-2 nucleocapsid (Sino Biological, Clone 05, 1:1000 dilution). Images were acquired using a Leica BOND III platform at the University of Virginia.

### Clinical sample collection for SARS-CoV-2 surveillance and variant analysis

Human clinical specimens for regional variant surveillance were collected between mid-2022 and early 2024 under IRB approval (Virginia Tech IRB# 20-852, Virginia Department of Health IRB# 70046) with informed consent. Nasopharyngeal swabs from suspected COVID-19 cases were submitted to Virginia Tech Schiffert Health Center’s Molecular Diagnostics Laboratory. A total of 14,446 were collected and 654 SARS-CoV-2 positive specimens (Cq ≤34) underwent whole genome sequencing. cDNA was synthesized using SuperScript IV Reverse Transcriptase, amplified with ARTIC nCoV-2019 primers (V4.1), barcoded using plexWell™ 384 Library Preparation Kit, and sequenced on MiSeq (Illumina). Bioinformatics analysis used containerized workflows including Trimmomatic trimming, Minimap2 alignment to MN908947.3 reference, Samtools processing, iVAR consensus generation, and Pangolin variant classification. Lineages were categorized as Omicron, XBB, JN.1, or other variants for temporal analysis. All sequences were deposited in GISAID and NCBI SARS-CoV-2 repositories.

### Detection of SARS-CoV-2 antibodies in biological fluids

Neutralizing antibodies were detected using multiple approaches: (1) qualitative lateral flow immunochromatography using the GenScript SARS-CoV-2 Neutralizing Antibody Detection Kit, ADEXUSDx COVID-19 Test, and NOVODIAX CoNAb SARS-CoV-2 Neutralizing Antibody Test for field testing; and (2) quantitative ELISA using Abcam COVID-19 S-Protein (S1RBD) Human IgG ELISA Kit. For species-independent detection by ELISA, samples from both human subjects (h.s.#1 collected prior to COVID infection in 05/2023, h.s.#1 collected post-COVID infection in 11/2023, and h.s.#2) and deer (A04, A07, A08, A09, A14, A23, and L-D4) were diluted appropriately and incubated with recombinant protein A/G (Thermo Scientific) to detect IgG from both species, followed by HRP-conjugated streptavidin detection. Absorbance was measured at 450 nm and concentrations were determined using calibration curves. Results were considered positive above 15 U/mL according to the manufacturer, who reports assay sensitivity of 94.74%, specificity of 98.57%, and overall accuracy of 97.75%.

### Viral RNA detection and stability testing

Samples collected from WTD and from humans (nasal swabs and milk) were analyzed for SARS-CoV-2 using RT-qPCR as described [34]. Transcription-mediated amplification using the Hologic Panther System was used on-site for a subset of WTDs in 2023 following the manufacturer’s instruction.

For stability assays, skim milk samples of human or WTD origin were spiked with ∼200,000 copies of heat-inactivated viral variants (A, B.1.1.7, BA.1.1.529) obtained from Zeptometrix. Samples were incubated at room temperature (∼22°C), 37°C, or 63°C for up to 60 min with aliquots collected in triplicate at specified timepoints. RNA was extracted using TRIzol and analyzed by RT-qPCR with quantification based on standard curves using 2019-nCoV_N positive controls (Integrated DNA Technologies).

### pH and ketone body determination

Milk pH was measured using colorimetric indicator strips (JT Baker) with detection range pH 4.5-10.0. Strips were wetted with 10-20 μL milk and compared to manufacturer reference scales with results recorded to the nearest 0.5 unit. Ketone bodies were detected using PrimeScreen Ketone Test Strips according to manufacturer instructions, with human urine from a consenting ketotic donor used as positive control.

### Quantitative determination of trace metals

Trace metal concentrations were determined by inductively coupled plasma mass spectrometry (ICP-MS) using an Agilent 7900 instrument. Milk samples were centrifuged at 12,000×g for 10 min at 4°C, and metabolites were extracted using methanol:water (4:1 v/v). After incubation at -20°C for 30 min and re-centrifugation at 14,000×g, supernatants were dried under nitrogen and reconstituted in 50% acetonitrile with 0.1% formic acid. Data acquisition and analysis were carried out using the Agilent 7900 ICP-MS instrument (Analytical Chemistry Research Core Laboratory, Virginia Tech) with Agilent MassHunter software. Peak intensities were normalized across samples and used for downstream comparative analysis.

### Total protease activity assay

Protease activity was measured in human and WTD milk specimens at room temperature using the Amplite® Universal Fluorimetric Protease Activity Assay Kit Green Fluorescence (AAT Bioquest), following manufacturer’s protocol A. The protease substrate was diluted 1:100 in 2x assay buffer to prepare the working solutions at pH 3, pH 7.6, and pH 8. Fluorescence intensity was measured at excitation/emission wavelengths of 490/525 nm using a microplate fluorescence reader (Molecular Devices).

### Multi-enzyme activity profiling

Human and WTD whole milk specimens were treated at either room temperature (∼22°C) or 100°C for 5 min, then placed on ice for 15 min prior to assaying various enzymatic activities using the API® ZYM system (BioMérieux). Temperature-treated samples were diluted 1:5 and incubated at 37°C for 4 h, followed by the addition of ZYM A and ZYM B reagents for color development. After a 5 min incubation, reactions were recorded using the manufacturer’s reference reading table, with intensity scored as “-“ (negative), and from weakest to strongest as “+,” “++,” “+++.”

### Lactoperoxidase (LPO) activity assay

Human and WTD milk specimens were treated at either room temperature (∼22°C) or 100°C for 5 min, then placed on ice for 15 min prior to assaying LPO using a Peroxidase Activity Assay Kit (Abcam). Reaction mixtures consisting of skim milk, OxiRed™ probe, and H₂O₂ substrate were incubated at 37°C for 3 min following manufacturer’s instructions, and the initial fluorescence (Ex/Em at 535/587 nm; A_0_) was recorded using a microplate reader (Molecular Devices). After a further 60 min incubation at 37°C, final fluorescence (A_1_) was measured. Peroxidase activity was calculated from a hydrogen peroxide standard curve, using the change in fluorescence [ΔA = A₁− A₀), and expressed as U/ul, where one unit corresponds to the oxidation of 1.0 μmol of H₂O₂ per minute at 37°C.

### Lysozyme activity assay

Lysozyme activity in skim milk samples was measured using the Lysozyme Activity Assay Kit (Abcam), following manufacturer’s instructions. Briefly, a substrate mixture prepared by combining lysozyme substrate with assay buffer was added to each sample and positive control wells. Plates were incubated at 37°C for 30 min in the dark. Fluorescence was measured immediately after stopping the reaction at Ex/Em = 360/445 nm using a FilterMax™ F3 Multi-Mode Microplate Reader. Lysozyme activity (mU/ml) was calculated using a 4-methylumbelliferone (4-MU) standard curve. One unit of activity is defined as the amount of enzyme that generates 1.0 μmol of 4-MU per minute at pH 5.0 and 37°C.

### Lactoferrin activity assay

Lactoferrin concentrations in WTD and human milk were quantified using the Bovine Lactoferrin ELISA Kit (Abcam) and the Human Lactoferrin ELISA Kit (colorimetric, Novus Biologicals), respectively, following manufacturers’ instructions. In both cases, skim milk was treated at either room temperature (∼22°C) or 100°C as previously described. Absorbance was measured at 450 nm immediately after stopping the reaction using a microplate reader (Molecular Devices SpectraMax X190), and lactoferrin concentrations were interpolated from the standard curve and adjusted for dilution.

### Functional protease inhibition and ions test studies

Human skim milk samples were incubated either at room temperature (∼22°C) for 20 min or at 100°C for 5 min, followed by 15 min on ice, in the presence or absence of an ion cocktail designed to mimic concentrations identified by ICP-MS in WTD milk. Protease inhibitors (final concentration: AEBSF 1 mM, aprotinin 800 nM, bestatin 50 μM, E64 15 μM, leupeptin 20 μM, pepstatin A 10 μM) were then added, or omitted (controls), and the samples were incubated at room temperature for an additional 5 min. Next, approximately 200,000 copies of SAR-CoV-2 A, B.1.1.7/Alpha, or B.1.529/Omicron (Zeptometrix) were spiked into the corresponding samples. Three equal-volume aliquots (45 µl) were withdrawn per treatment, and viral RNA was purified and quantified as described in [34]. Comparable protease inhibitor experiments were carried out using WTD milk, except that ions were not supplemented in this case.

### Milk reconstitution assays

Human milk was supplemented with ions to mimic deer milk concentrations (Na⁺ 20.91 mM, Mg²⁺ 2 mM, P 8.6 mM, K⁺ 20.8 mM, Ca²⁺ 0.74 mM, Fe³⁺ 6.7 μM, Zn²⁺ 11 μM) and/or LPO (0.4 U/μl, Sigma) to assess their individual and combined effects on viral RNA stability. Approximately 200,000 viral RNA copies were added to treated samples, incubated for 5 min at room temperature, then processed for RT-qPCR analysis.

### Statistical analysis

All experiments were performed in triplicate unless otherwise noted. Data are presented as mean ± standard deviation (SD) as indicated. Statistical analyses included appropriate controls and standardization to input viral loads for comparative studies. RT-qPCR quantification was based on standard curves using certified reference materials. Differences between two groups were analyzed using the Student’s t test. Differences between more than two groups were analyzed by ordinary one-way analysis of variance (ANOVA) followed by a Holm-Sidak’s multiple comparisons test. All statistical analyses were performed with GraphPad Prism7 (Software Inc.), and p values of 0.05 or less were considered statistically significant.

### Ethics Statement

All human sample collection and research activities were reviewed and approved by the Institutional Review Board at Virginia Tech (IRB#20-852 and IRB#24-474) and by the Virginia Department of Health (IRB#70046). Written informed consent was obtained from all participants prior to enrollment.

For white-tailed deer (WTD) samples, the Virginia Department of Wildlife Resources was consulted prior to the legal harvest of game animals. Samples were obtained opportunistically post mortem from legally harvested wildlife; therefore, the study did not require Institutional Animal Care and Use Committee (IACUC) oversight. The work was reviewed and determined to be exempt under Category 2 and Category 4 by the Centra Health Institutional Review Board (CHIRB 0601e).

## Acknowledgments

We thank Dr. J. Webster for comments and proofreading of the manuscript, all members from the Molecular Diagnostics Laboratory, Fralin Biomedical Research Institute for feedback on the manuscript. We also wish to thank the hunters and private landowners for the harvest of WTDs utilized in this study, and the staff at Pathology Consultants of Central Virginia.

This study was funded by the USDA-APHIS and the Fralin Biomedical Research Institute to C.V.F. Additional funding was provided by Pathology Consultants of Central Virginia and a research stipend from the University of Virginia Department of Pathology to A.B. The funder played no role in study design, data collection, analysis and interpretation of data, or the writing of this manuscript.

## Author Contributions

H.A.R. and D.S.R. contributed to specimen acquisition and to the data presented in Figures 1B–C. A.P.B. and D.S.R. performed histopathological analyses and contributed to Figure 1G. L.T. contributed to Figures 2–5. W.S. conducted laboratory analyses and contributed to the data presented in Figures 1D–E, 2, 3A, 3E, and 5B. A.C., K.B., M.G.U., C.R., D.M., and D.G.S.C. contributed to research methodology, genomic data analysis, and data interpretation across all figures. I.A. performed all statistical analyses presented in the study. All authors provided critical feedback on manuscript drafts. L.T. prepared the initial full manuscript draft, and D.S.R. contributed specific sections. D.S.R. and C.V.F. designed the study and provided project administration throughout the research. C.V.F. prepared the final version of the manuscript.

## Supporting Information

**Figure S1.** Geographic distribution and hunting context of WTD specimen collection in central Virginia, USA. WTD harvest occurred on private land with permission in central Virginia during the 2022-2024 hunting seasons in Amherst, Bedford, and Nelson counties. Participating licensed hunters preferentially targeted does (approximately 66% of total harvested WTD) at landowner request, resulting in 8 lactating does (approximately 25% of harvested does) utilized for this study. The map shows county boundaries with color-coding corresponding to sample density. Individual sampling locations are marked with colored dots: orange dots indicate female deer. The lactational WTD county distribution table (lower right) details the geographic harvest locations of the 8 lactating does. The inset map (left) shows the geographic location within Virginia, USA. The 2022-2024 central Virginia WTD harvest table (upper right) provides county-specific information including total area (hectares) and three-year estimates (mean) for total number, density, and percent female of reported WTD harvest based on Virginia Department of Wildlife Resources (VDWR) data. Note the relatively high WTD density, especially in Bedford County and to a lesser extent in Amherst and Nelson counties. This hunting approach reflects VDWR management goals to stabilize county-specific populations through liberalized regulated hunting, especially for females, to minimize impacts on herd health, property, and human safety (http://www.virginiawildlife.gov/wildlife/deer/management-plan).

## Notes

### Competing Interest Statement

The authors have declared no competing interest.

